# Osteopontin in colitis-associated carcinoma

**DOI:** 10.1101/2023.12.15.571952

**Authors:** Subhakankha Manna, Maximilian Sehn, Danielle Cardoso da Silva, Jakob J. Wiese, Claudia Heldt, Izabela Plumbom, Thomas Conrad, Cora C. Husemann, Karsten Kleo, Lorena Derêzanin, Violaine Dony, Saeed K. Farahani, Franziska Weiss, Federica Branchi, Anja A. Kühl, Simon Schallenberg, January Weiner, Sefer Elezkurtaj, Benjamin Weixler, Jörn Gröne, Britta Siegmund, Michael Hummel, Michael Schumann

**Author notes:** equally contributing (Subhakankha Manna, Maximilian Sehn, Danielle Cardoso da Silva).

## Abstract

Patients with ulcerative colitis (UC) and Crohn’s disease (CD) face a lifelong risk of developing colitis-associated carcinoma (CAC). Current insights into CAC development originate from murine CAC models, while human CAC studies primarily focus on mutational analysis of patient samples. Although the mutational landscape reveals distinct patterns and frequencies compared to colorectal cancer, it falls short in elucidating the inflammatory mechanisms of CAC development. Consequently, we adopted a multi-omics approach to unravel CAC carcinogenesis from an immunological perspective. Our data revealed a robust upregulation of *SPP1* gene in CAC at both RNA and protein levels, expressed by *CD68+* macrophages. In vitro OPN stimulation demonstrated no direct effect on intestinal epithelial organoids. However, the mutually exclusive spatial location of *SPP1/OPN+* macrophages and *CD8+* T cells suggests a crucial indirect role of *SPP1/OPN* in mediating an immunosuppressive tumour microenvironment.

## Introduction

About 0.4% of Europeans suffer from one of the two inflammatory bowel diseases, Crohn’s disease (CD) or ulcerative colitis (UC). Besides the significant reduction in quality of daily life in active CD or UC secondary to chronic diarrhea, hematochezia, episodes of severe abdominal pain and reduced physical fitness, these diseases pose an increased risk to patients to develop colitis-associated carcinoma (CAC) ^1, 2^. In numbers, CAC accounts for approx. 10-15% of deaths in IBD patients ^3^. In a study that exploited carcinoma findings in UC surveillance colonoscopies, the cumulative incidence of CAC was described to rise from 2.5% at 20 years to 7.6% at 30 years of colitis duration ^4^. The advent of targeted therapies using biologicals has contributed to a reduction of CAC occurrence, pointing to the role of mucosal inflammation in colitis-associated carcinogenesis ^3^. Nevertheless, a more recent meta-analysis identified the relative risk of patients suffering from inflammatory bowel disease to develop CAC to be 2.4-fold higher compared to the risk of the general population developing sporadic colorectal carcinoma (CRC) ^5^.

Differences in clinical presentation as rapid progression, presentation of CACs as interval carcinomas and occurrence of gene mutations distinct from sporadic CRC point to a CAC-specific tumor biology. As such, APC mutations, which are a cause for nuclear b-catenin translocation and consecutive stimulation of cell growth and inhibition of apoptosis, occur not as an initial step as found in sporadic CRCs but as a late carcinogenetic event ^6, 7^. Moreover, mutations of the TP53 gene occur early on in CAC, prolong the activation of NF-kB signaling, aggravate the extent of colitis and accelerate the progression to high grade dysplasia and invasive carcinoma while TP53 is mutated only as a rather late step in sporadic CRCs ^7, 8^. Another significant discrepancy to CRCs involves the role of pro-inflammatory cytokines as interleukin-6 (IL-6), IL-11, IL-22 and TNF-α ^9-11^. Signaling of these cytokines occurs through kinase cascades including JAK/STAT, MAPK, PI-3 kinase and regulates expression of a multitude of putative tumor-relevant target genes including cytokine and cytokine receptor genes as IL-21, IL-21R and IL-23R and further transcription factors as RORγT, IRF4, and Batf ^12^. Moreover, it was shown, that IL-22 and its natural counterpart IL-22 binding protein are directly involved in murine colon carcinogenesis ^13^. One mechanism, by which cytokines as IL-22 or TGF-b may contribute to carcinoma development is epithelial-to-mesenchymal transformation (EMT) ^14, 15^. In this study, we identify another cytokine, osteopontin (OPN), to be highly expressed in inflammation-associated carcinogenesis in IBD patients.

## Results

### Clinical characteristics of the patients included in the study

To better understand CAC pathogenesis, patients who underwent surgery and were diagnosed with Crohn’s disease-colitis, ulcerative colitis, colitis-associated colon cancer or sporadic colorectal cancer were retrospectively recruited from the Charité database after approval by the Charité ethical commission (EA/1/204/14). Inflammation-free borders of colon resections of diverticulitis patients were selected as the control population. CAC patients were categorized according to the previous IBD condition of the patients in Crohn’s disease-associated CAC (CDAC) and ulcerative colitis-associated CAC (UCAC). In total, 20 CAC’s were included, that were in the majority of the mucinous type on histologic examination (**Table 1**). Of note, the inflammatory colitis activity in patients with IBD was higher when compared to the CAC patients, as the surgery in these patients was performed for their IBD, while the need for surgery in the CAC patients resulted from the carcinoma (**Table 1**).

**Table 1:**
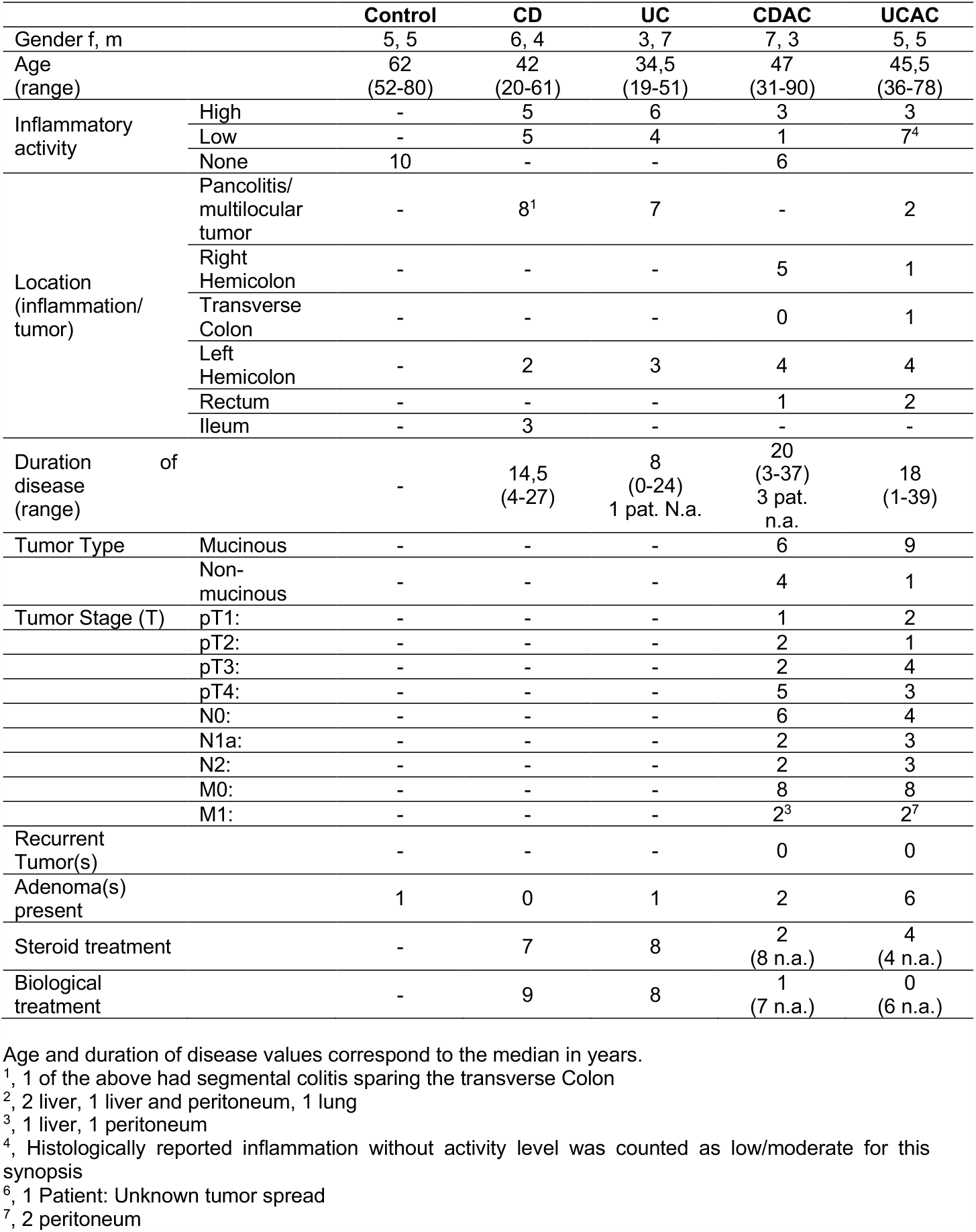
Clinicopathological characteristics.

### Differential gene expression in colitis-associated carcinoma compared to colitis

To have a comprehensive view on differential gene expression in CAC, we compared the gene expression in colon tissues of patients with an IBD-associated colon carcinoma to either Crohn’s colitis or UC surgical colon samples. For that, a Nanostring analysis was performed examining 654 genes (624 genes from the human immunology panel plus 30 custom genes). SPP1 gene was robustly upregulated in both UCCAC and CDCAC **(Figure 1B)**. Heatmaps of the 20 most upregulated and 20 most downregulated genes for CDAC vs. CD and UCAC vs. UC showed that CD and CDAC patients clustered separate from each other (**Figure 1C**) and that the clustering was almost perfect for UC and UCAC patients except for one UC patient (**Figure 1C**). The differentially regulated genes are listed in **Tables 2 and 3**. SPP1, the gene for osteopontin (OPN), was found to be the most upregulated gene in both comparisons, even though the two patient groups analyzed were totally independent from each other. Other significantly regulated genes included genes of cell signaling pathways (MAPK/ERK, DUSP4, SRC), chemokine genes and B-cell genes. Gene set analysis revealed genes for the adaptive immune system, lymphocyte activation and chemokine signaling to be mostly downregulated (**Table 4**).

**Table 2:** 10 most up- and down-regulated genes in CDAC *vs*. CD.

|  | **Gene** | **Fold change** | **p-value** |
| --- | --- | --- | --- |
| **Up-regulated genes** | SPP1 | 18,29 | <0,001 |
|  | FN1 | 6,68 | <0,001 |
|  | DUSP4 | 4,60 | <0,001 |
|  | CLDN2 | 4,40 | 0,034 |
|  | CCL26 | 3,42 | <0,001 |
|  | CD276 | 3,01 | <0,001 |
|  | THY1 | 2,83 | 0,009 |
|  | ICAM5 | 2,78 | 0,010 |
|  | F2RL2 | 2,77 | 0,010 |
|  | CLEC5A | 2,74 | 0,012 |
| **Down-regulated genes** | PIGR | -23,69 | <0,001 |
|  | CXCL13 | -11,11 | <0,001 |
|  | MS4A1 | -9,77 | <0,001 |
|  | CD79A | -7,51 | <0,001 |
|  | NOS2 | -7,34 | 0,002 |
|  | CR2 | -6,58 | <0,001 |
|  | CCL19 | -6,44 | <0,001 |
|  | PTPRC (RA) | -6,33 | <0,001 |
|  | TNFRSF17 | -6,12 | <0,001 |
|  | TNFRSF13B | -5,38 | <0,001 |

**Table 3:** 10 most up- and down-regulated genes in UCAC *vs*. UC.

|  | Gene | Fold change | p-value |
| --- | --- | --- | --- |
| **Up-regulated genes** | SPP1 | 8,41 | <0,001 |
|  | CCL26 | 2,91 | 0,022 |
|  | GRHL2 | 2,71 | <0,001 |
|  | DUSP4 | 2,67 | 0,030 |
|  | CCND1 | 2,65 | 0,014 |
|  | FN1 | 2,22 | 0,039 |
|  | CEACAM6 | 2,05 | 0,032 |
|  | CD9 | 1,78 | 0,013 |
|  | SRC | 1,70 | 0,011 |
|  | TRAF2 | 1,68 | 0,008 |
| **Down-regulated genes** | DEFB4A | -18,33 | <0,001 |
|  | S100A8 | -14,97 | <0,001 |
|  | PIGR | -11,10 | 0,011 |
|  | NOS2 | -10,15 | <0,001 |
|  | S100A9 | -8,56 | <0,001 |
|  | MS4A1 | -8,20 | <0,001 |
|  | IRF4 | -7,90 | <0,001 |
|  | CXCL13 | -7,31 | 0,001 |
|  | TNFRSF13B | -7,26 | 0,001 |
|  | IL1B | -7,00 | <0,001 |

**Table 4:** Analysis of gene sets.

|  | No of genes included | Up- (↑) / down- (↓) regulated genes | Directed significance score |
| --- | --- | --- | --- |
| MHC II antigen presentation | 14 | 5↓ | -3.18 |
| Adaptive immune system | 137 | 39↓ | -3.12 |
| Lymphocyte activation | 241 | 3↑ 62↓ | -3.07 |
| TNF family signaling | 49 | 7↓ | -2.74 |
| Cell adhesion | 56 | 2↑ 17↓ | -2.57 |
| Cytokine signaling | 259 | 3↑ 44↓ | -2.55 |
| NF-κB signaling | 62 | 10↓ | -2.53 |

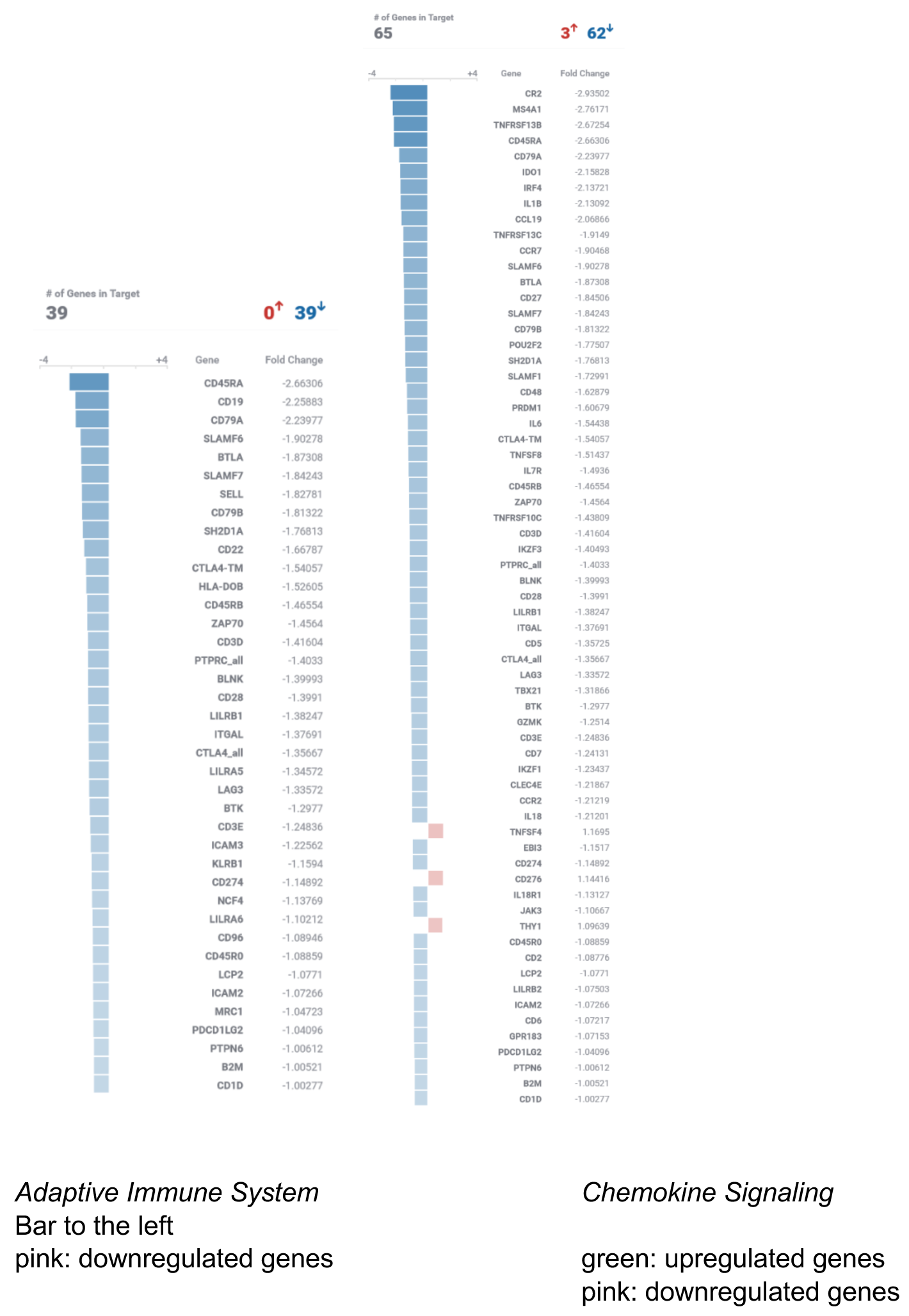

**Figure 1.**
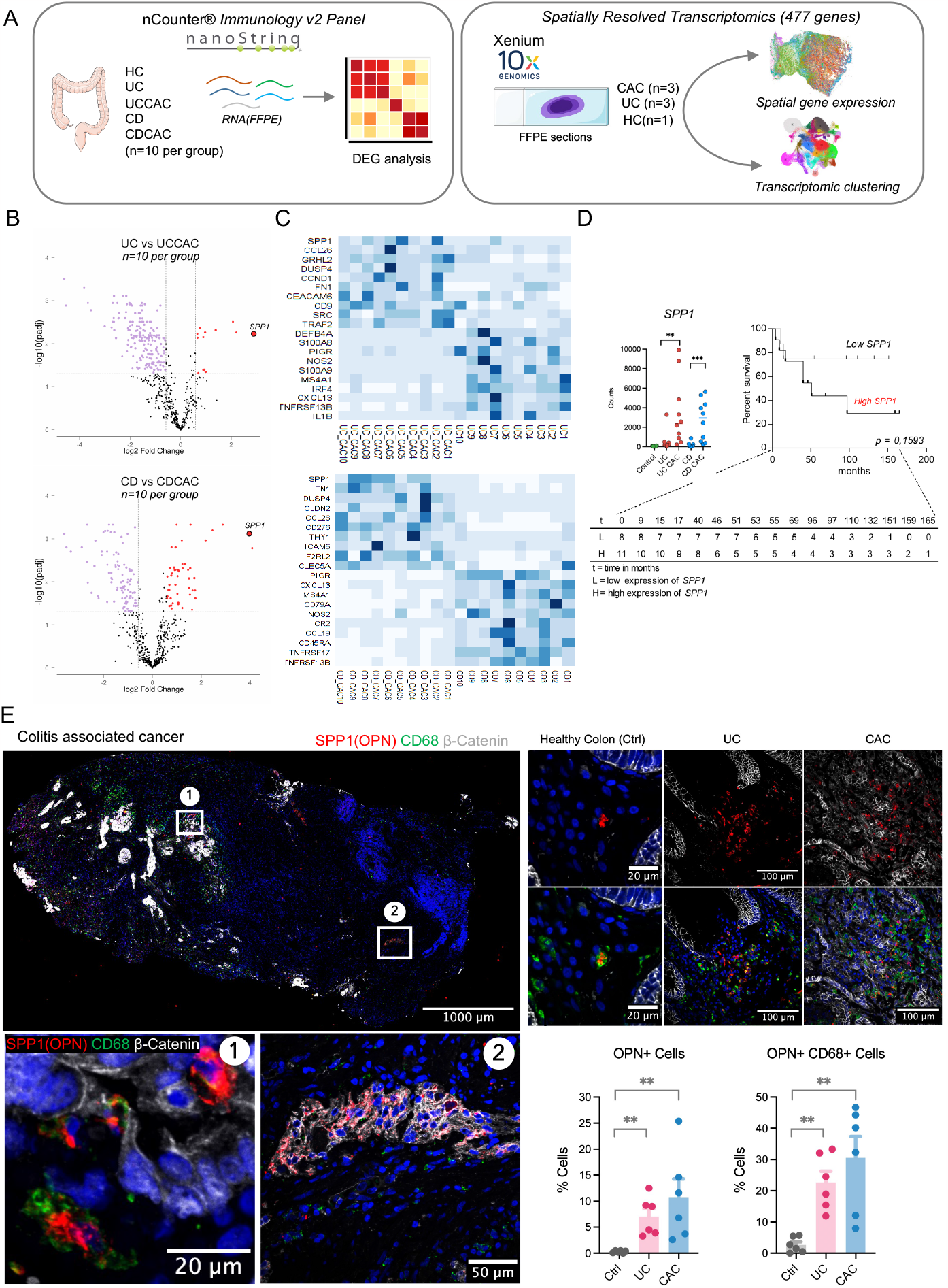
A. Study design. B Nanostring Immune Panel v2 Volcano plots of UC vs UCCAC and CD v CDCAC reveals SPP1 to be upregulated in both cohorts. C, Heatmaps with top 20 genes, D. KM plot using SPP1, E. Heterogenous OPN expression in CAC, F. Quantification of OPN expression across Control, UC and CAC patients (n=3).

### Expression of OPN, its cellular source and disease prognosis

Since OPN was independently found as the most up-regulated gene in both comparisons, we decided to further investigate it. Analysis of the OPN counts from the Nanostring evaluation revealed a significant scatter, especially for the CAC conditions, with two separate groups of patients expressing either high or low OPN (**Figure 1C**). Stratifying patients along these groups (i.e., high or low OPN expressers) showed a tendency for a lower survival rate of the high OPN-expressing patients (**Figure 1D**), even though this correlation did not reach statistical significance. Analysis of OPN protein levels by a multi-staining immunehistological approach confirmed not only the strong upregulation in CAC samples but revealed furthermore a significant inhomogeneity when larger areas of tumor tissues were examined by stitching large tumor sections (in average approx. 5000 µm in diameter) after spinning disk imaging (**Figure 1E**). The majority of OPN-expressing cells were identified to be monocytes by CD68 immunostaining and – to a significant lesser extent – myenteric plexus cells (**Figure 1E**). Spatial resolution of RNA expression (10x Genomics, Xenium platform) confirmed the strong SPP1 expression in CAC tissue and further subclassified OPN-expressing cells as *CD68+ macrophages* (**Figure 2A**), and identified a macrophage subset in CAC that are *SPP1+ APOE+ CD163+* (**Figure 2C, Supplementary Fig 1**). SPP1 clusters were consistent across CAC patients but not in UC patients (**Supplementary Fig 1)**. Moreover, the analysis of OPN expression by splice variant-specific probes allowed the identification of the prevailing OPN splice variant in CACs to be SPP1v1, while it was SPP1v2 in UC tissue (**Figure 3A-C**).

**Figure 2.**
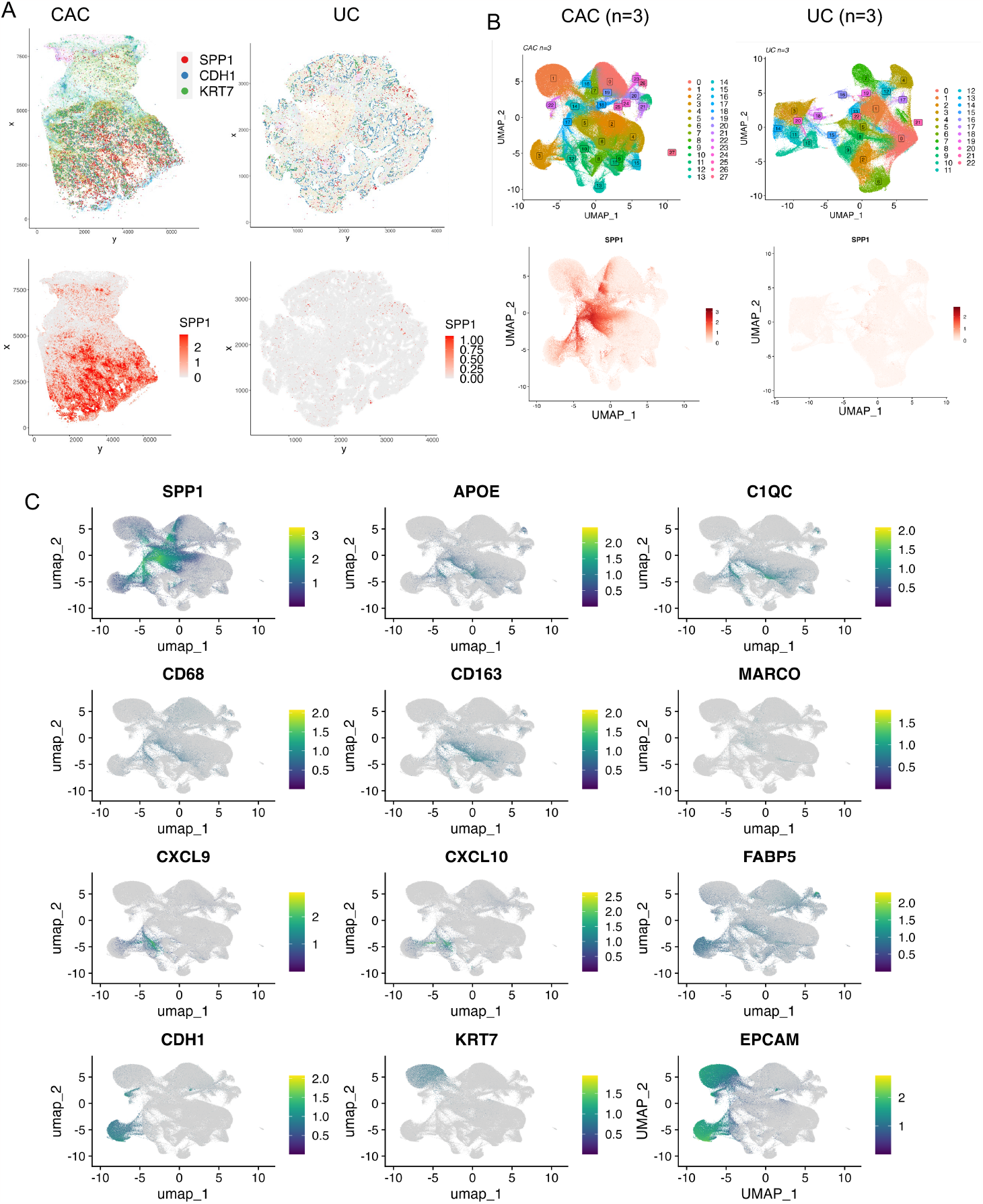
Spatial heterogeneity of SPP1 expression in CAC. (A) Spatial gene expression plots of SPP1, CDH1 and KRT7 expression in CAC and UC (B) Unsupervised clustering results of CAC(n=3) and UC(n=3). (C) UMAP plots of genes in SPP1 macrophage cluster.

**Figure 3.**
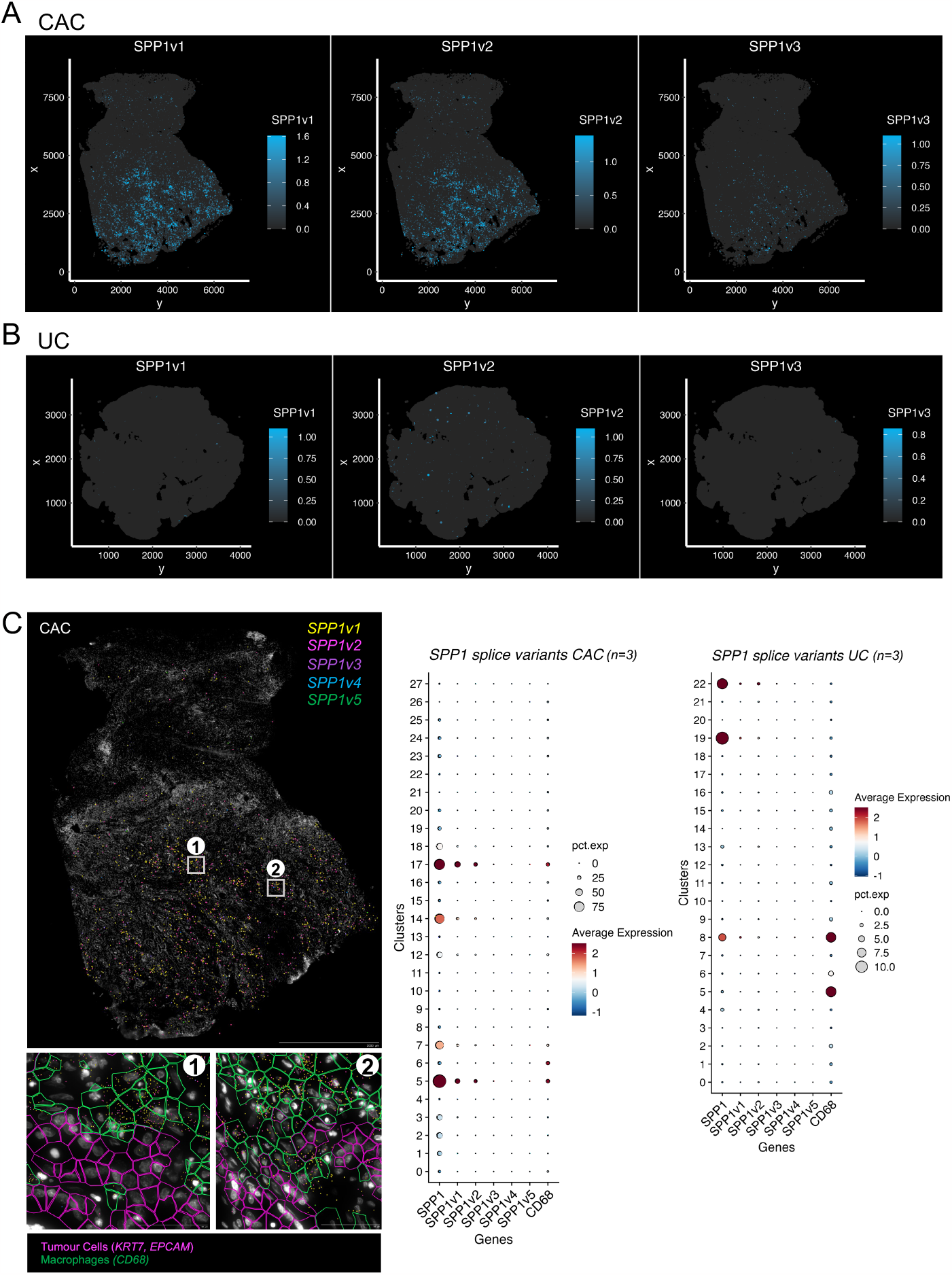
SPP1 splice variants in CAC and UC. A. SPP1v1, v2, v3 transcript density plots of CAC tissues, B. SPP1v1, v2, v3 transcript density plots of UC tissues, C. SPP1v1, v2, v3 transcript at single cell resolution in CAC, Dotplots showing the transcriptomic clusters and SPP1 splice variants in CAC and UC (n=3).

### Identification of putative OPN targets

Next, we sought to identify putative OPN targets by analyzing the expression of previously described OPN receptors in various cell types. This analysis comprised the previously as a receptor for hyaluronic acids described glycoprotein CD44 including its alternative splicing variants (CD44, CD44 variant 3, CD44 variant 6, CD44 variant 7) and various integrin α and b chains that had previously been described to transmit OPN signals (**Figure 4**). Of note, the spatial resolution of the Xenium-RNA expression analysis allowed to take relative locational proximities of the respective gene expressions into consideration. This analysis revealed that CD44 is expressed in multiple cell types in CAC and there was little to no expression of CD44 splice variants. CD44 immunostainings revealed its expression on OPN+ macrophages, indicating an autocrine loop. Xenium RNA expression identified ITGαV and ITGβ6 in CAC epithelial clusters samples (**Figure 4A-E**). Further exploration of top DEG’s in epithelial cluser revealed strong expression of Claudin 7. This tumour cluster showed strong expression of KRT7, EPCAM, CTNNB1 and GLUT1 **(Figure 5 A-B)**.

**Figure 4.**
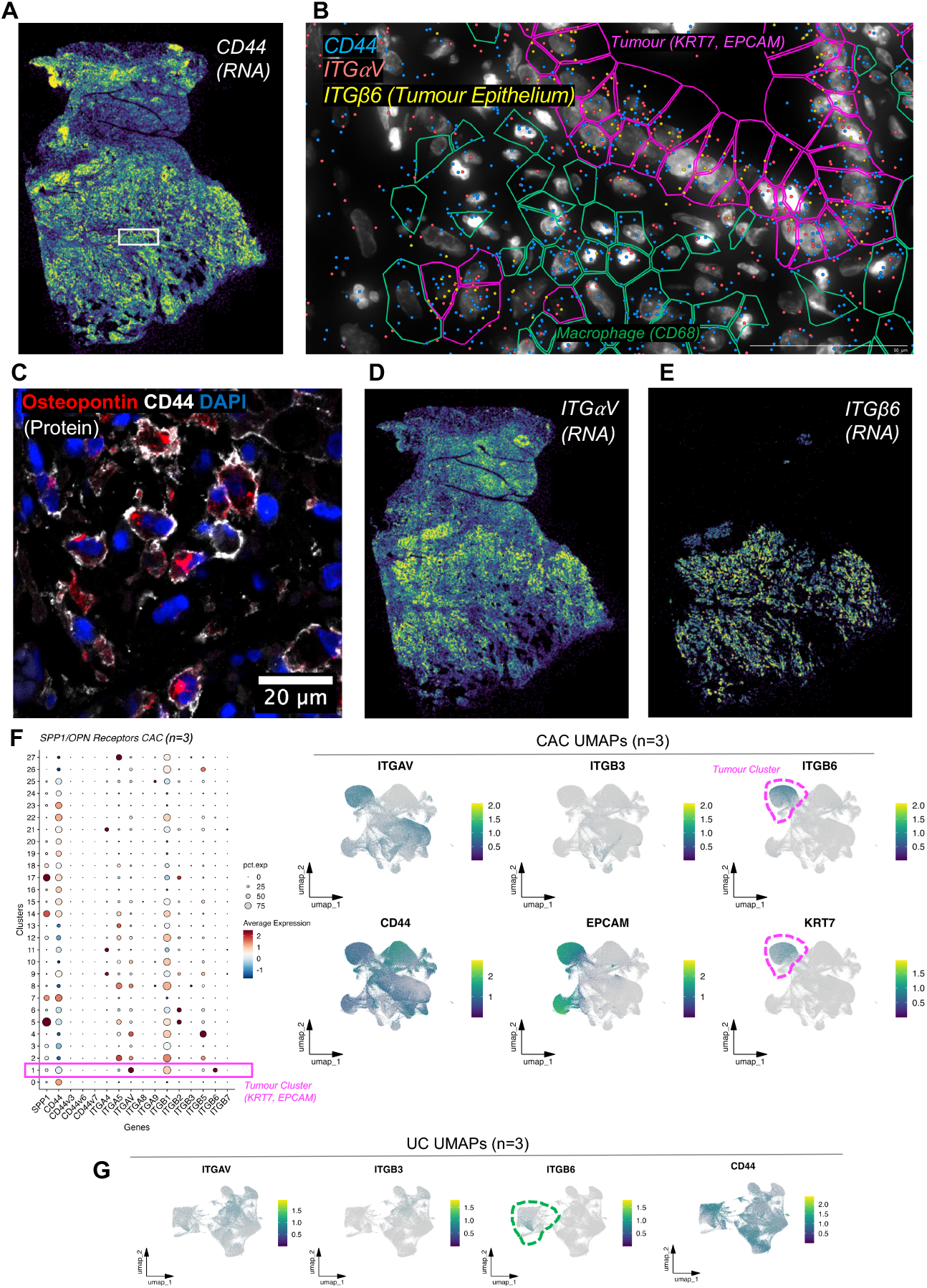
OPN Receptors in CAC. A, D, E. Density maps of CD44, Integrin aV and β6 expression in CAC B. CD44, Integrin aV and β6 transcripts at single cell resolution. F. Dotplots showing transcriptomic clusters in CAC and UMAPs of putative OPN receptors in UC and CAC (n=3).

**Figure 5.**
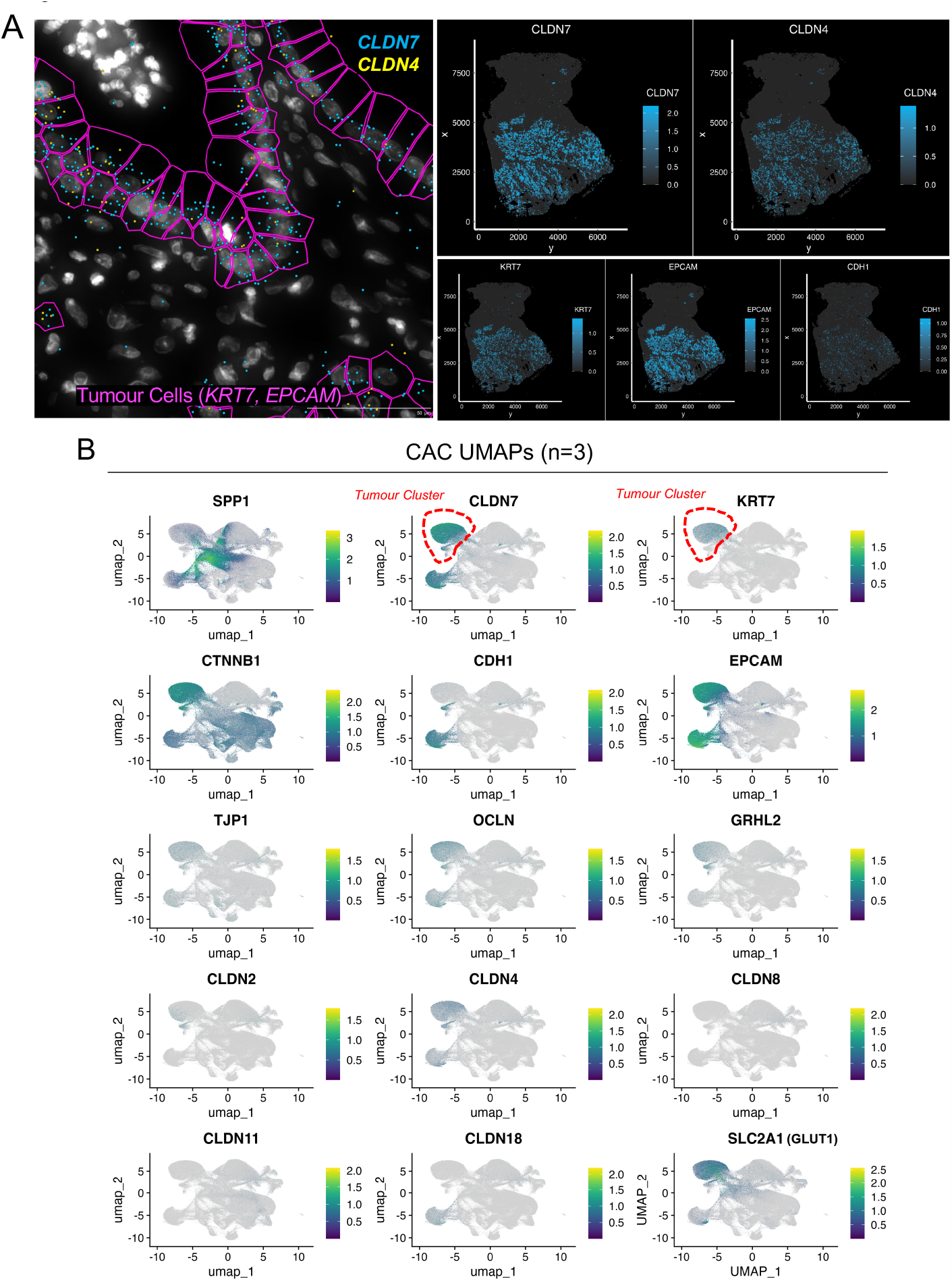
A. Tight Junction genes CLDN7 and CLDN 4 at a single cell resolution in CAC. B. UMAPs of TJ genes in CAC.

### Immunosuppressive Niche in CAC

We identified a skew in the tissue distribution of lymphocytes within CACs: Regions with strong OPN expression were devoid of CD8^+^ T-lymphocytes and vice versa (i.e. regions with low OPN expression were CD8^+^ lymphocyte-rich (**Figure 6A-C**). Together with the fact, that the gene set analysis of the Nanostring RNA expression had pointed to a downregulation of (CD8^+^ lymphocyte signaling) and (lymphocyte activation) and that others had shown a function for OPN in CD8^+^ lymphocyte activation in experimental murine colitis (Klement, 2018), we hypothesized, that OPN might suppress CD8^+^ lymphocyte-related tumor surveillance processes in CAC.

**Figure 6.**
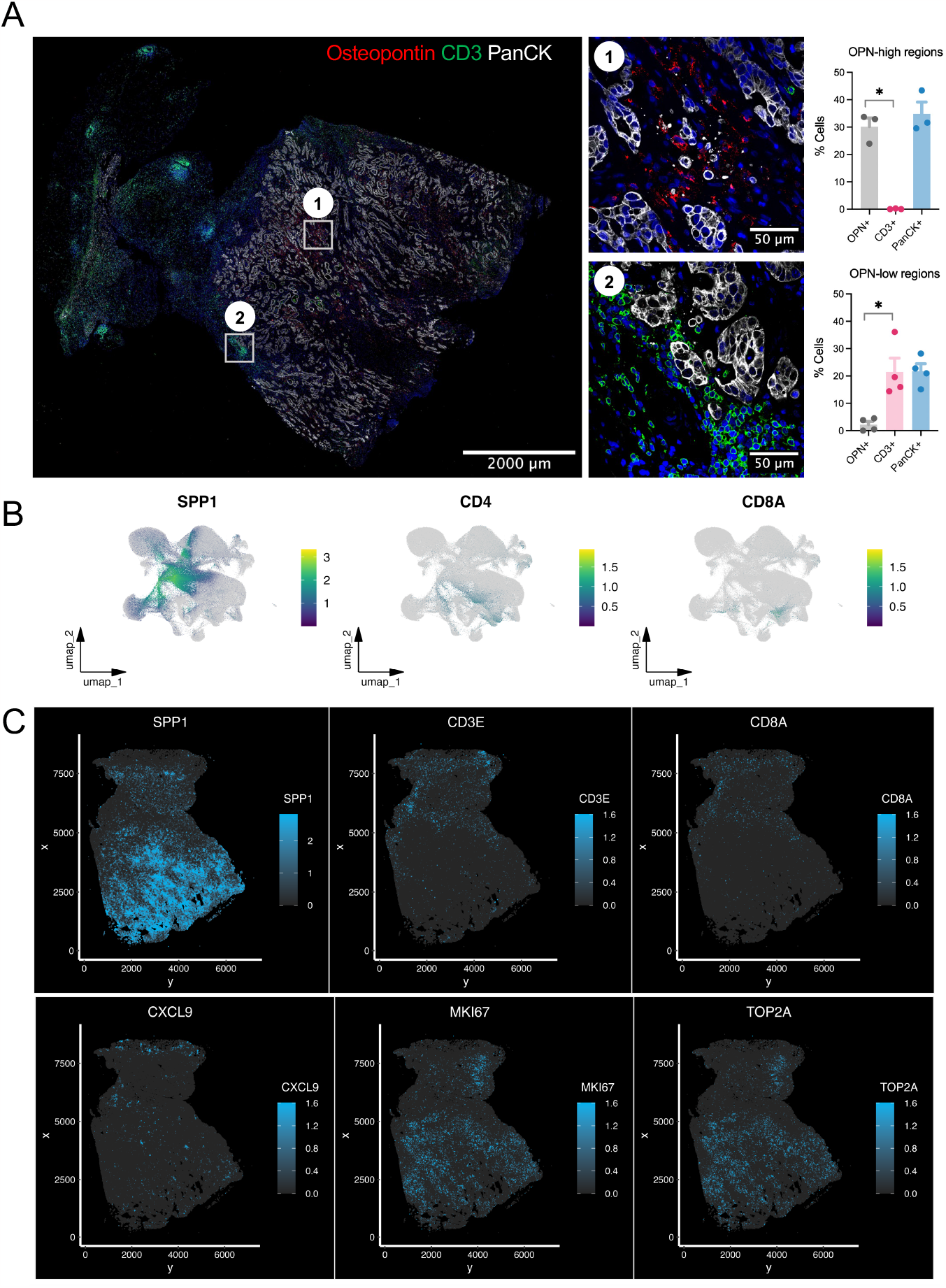
Immunosuppressive niche in CAC. A. Immunostainings of CAC tissues showing mutually exclusive expression of OPN and CD3. B. UMAP plots of CAC showing SPP1, CD4 and CD8A clusters. C. Transcript density maps of SPP1, CXCL9, CD3, CD8A and cell cycle genes.

### Epithelial-to-mesenchymal transformation as a putative mechanism for OPN action on CACs

It has been previously described for OPN, that it activates epithelial-to-mesenchymal transition (EMT) as one pathway having the potential to contribute to carcinogenesis. In our data set and in line with this assumption, the transcription factor ZEB1 and the extracellular matrix protein fibronectin (FN1) were amongst the most up-regulated genes (**Tables 2 and 3**). To further substantiate these findings, immunostainings were performed for EpCAM, E-cadherin (**Figure 7A-B**) and vimentin (data not shown). Interestingly, blinded analysis of the slides revealed a tendency for lower EpCAM and lower E-cadherin but higher expression of vimentin in CAC tissues, supporting the hypothesis that EMT might be an active process in CAC (**Figure 7A**).

**Figure 7.**
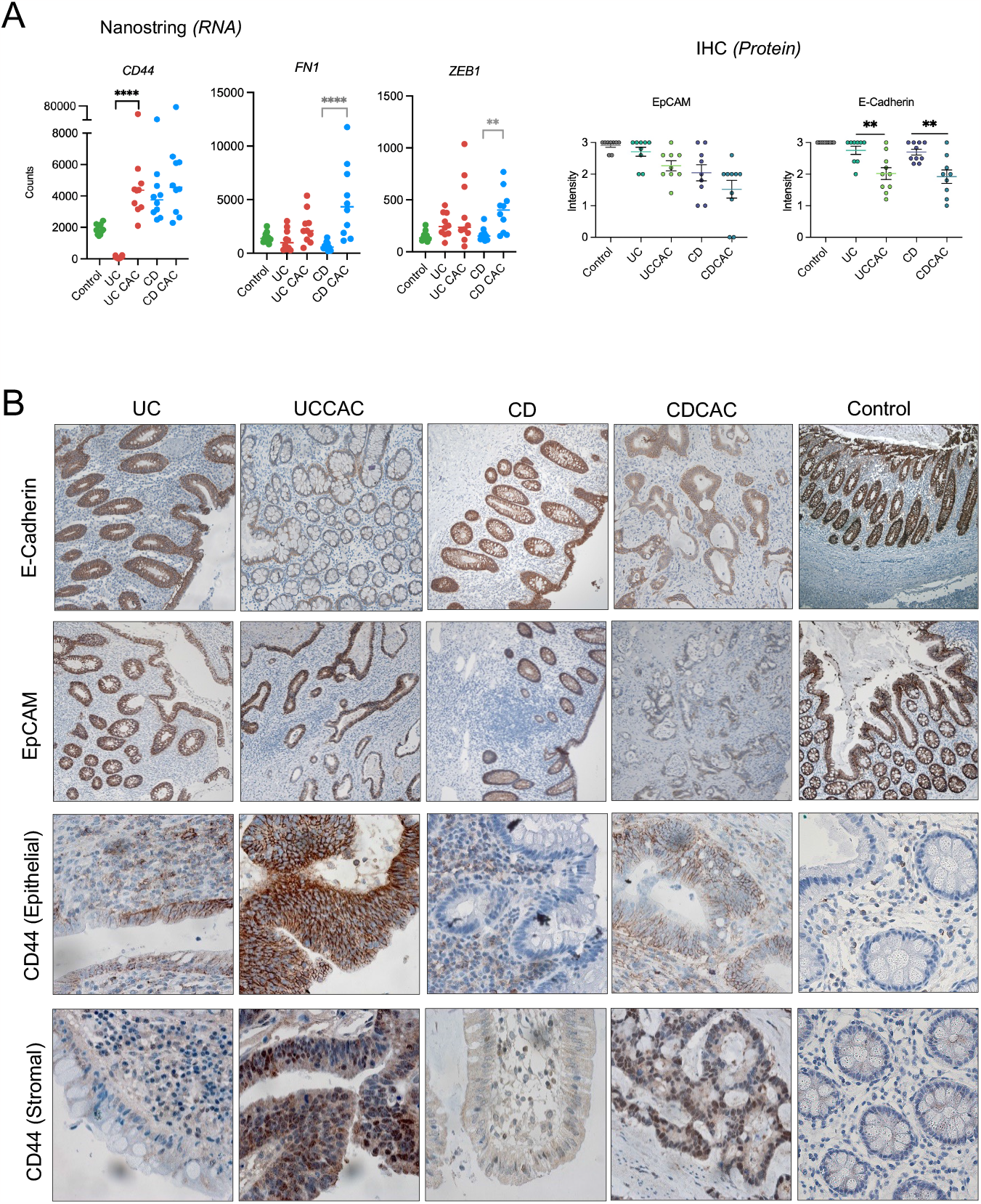
EMT in CAC. A. Nanostring gene expression dotplots showing EMT genes CD44, FN1 and ZEB1. B. IHC staining of epithelial proteins E-Cadherin, EpCAM and CD44 in UC, UCCAC, CD, CDCAC and Controls(n=10 per group)

To clarify, if OPN might have the potential to induce EMT in the process of colitis-associated carcinogenesis, we established these organoids from healthy donors, UC patients (Sato 2011). and CAC patients (Fujii 2018, Driehuis 2020). These organoids expressed high levels of putative OPN receptors including CD44 and Integrins when grown in stem cell medium. We then performed a series of RT-qPCR analyses to identify the effects of OPN, these gene panels included EMT genes, cell lineage genes and genes that were reported previously to be induced by OPN stimulus. However, these genes were not regulated by OPN stimulus and furthermore RNA sequencing analysis confirmed the following results (**Figure 8**) that these UC organoids do not respond to OPN, indicating an indirect role of OPN dependent tumorigenesis in CAC.

**Figure 8.**
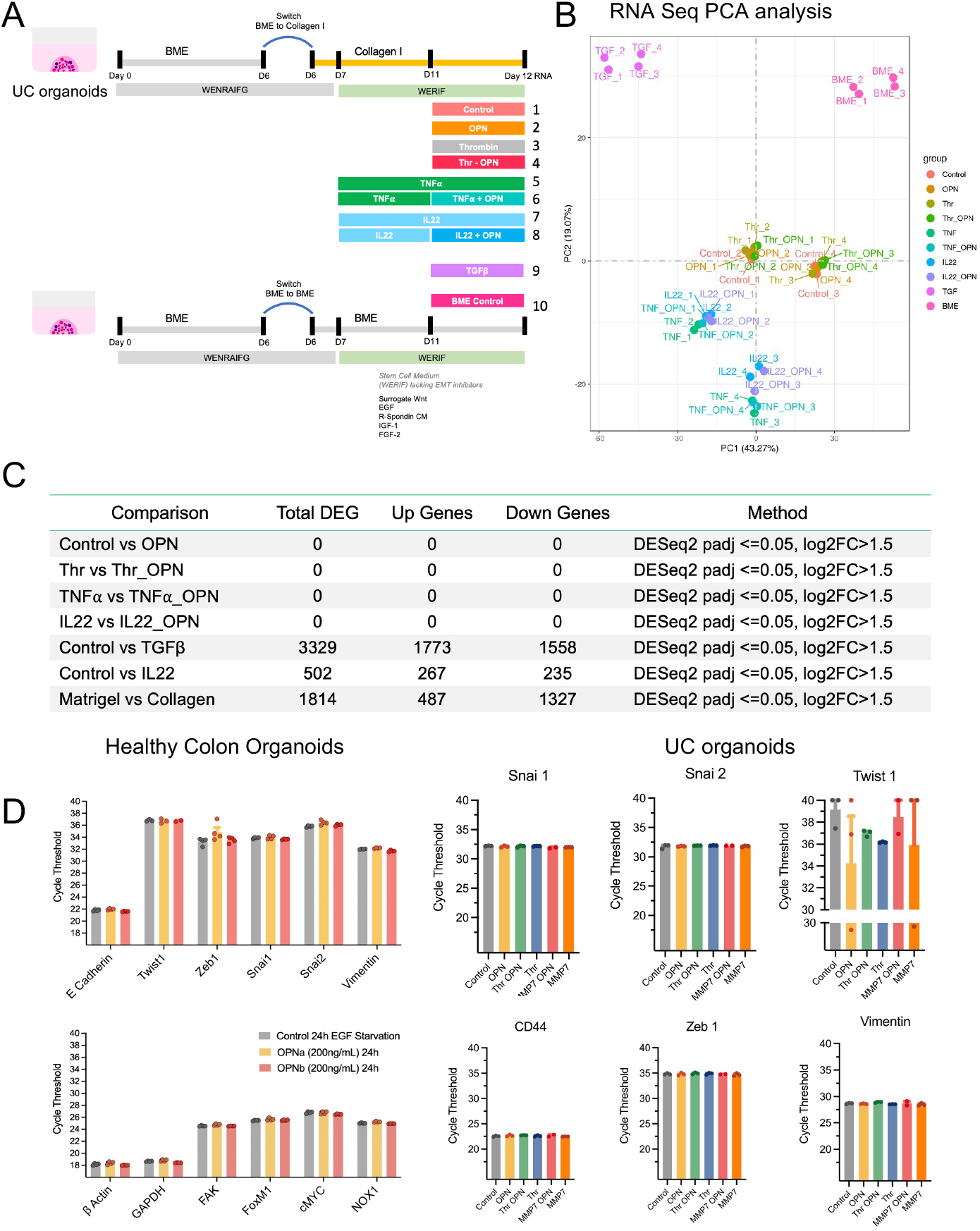
RNA Seq analysis of UC organoids. A. Experimental details of OPN stimulated UC organoids. B. PCA analysis of OPN stimulated UC organoids. C. Table of differentially expressed genes in UC organoids in response to OPN. D. RT-qPCR analysis EMT genes in colon organoid stimulated with OPN.

## Materials and Methods

### Nanostring Immunology v2 panel

We obtained FFPE specimens (Charité Biobank) from 40 patients that suffered from either one of the IBDs or CACs and furthermore inflammation-free healthy controls (10 patients per group). These samples were micro-dissected, followed by RNA isolation and Nanostring gene expression analysis using the Immunology v2 panel (630 genes).

### Spatially resolved transcriptomics

To spatially resolve expression of a set of 477 genes at a sub-cellular resolution, we analyzed surgical FFPE tissue sections of UC and UC-CAC patients using the Xenium platform from 10x Genomics. We used the Human Multi-tissue and Cancer panel as base and added 100 custom genes, these genes were selected on the basis of SPP1 biology and tumour microenvironment.

### Xenium sample preparation, imaging, and segmentation

Samples were processed according to manufacturer protocols “Xenium In Situ for FFPE – Tissue Preparation Guide CG000578 | Rev C”, “Probe Hybridization, Ligation & Amplification, User Guide CG0000582” and “Decoding & Imaging, User Guide CG000584. Briefly, 5 μm thick sections were prepared from FFPE tissue of UCCAC(n=3), UC(n=3 and HC(n=1). All 3 UCCAC tissue sections were placed on one slide and all 3 UC and 1 HC tissues were placed on the second slide. This was followed by deparaffinization & decrosslinking of the sections. Slides were then placed in xenium cassettes and padlock probes were incubated overnight, followed by rolling circle amplification, autofluorescence quenching and nuclear staining. Slides were added to the Xenium Instrument for FOV selection followed by the iterative cyclic imaging. Segmentation data and transcript level decoding data was generated on board using xenium-1.6.0.8 analysis software.

### Transcriptomic clustering of segmented cells

For downstream analysis, we used the package Seurat (v 5.0.1) (Stuart 2019) in R (v4.3.2). We followed the vignette from Satija Lab. (https://satijalab.org/seurat/articles/seurat5_spatial_vignette_2). Briefly, SCTransform was performed on all samples followed by filtering out cells with less than 20 genes. We then merged the CAC (n=3) gene matrices and UC (n=3) gene matrices into seurat objects. SCT was set as default assay followed by computing principal components (PCs) in 30 dimensions. We then selected these PC’s and used FindNeighbour and RunUMAP functions. We then portioned the shared nearest neighbour (SNN) graph using a resolution of 0.5 and identified 28 clusters in UCCAC and 23 clusters in UC. To identify the DEG in each cluster, we used PrepSCTFindmarkers function.

### DNA extraction

gDNA extraction from formalin-fixed paraffin-embedded (FFPE) tissue sections was carried out using Maxwell 16 FFPE plus LEV DNA purification kit (AS1720; Promega, Madison, WI, USA) according to the manufacturer’s protocol. Subsequently, fluorescence-based quantification was performed using the Qubit dsDNA HS assay kit (Thermo Fisher Scientific, Waltham, MA, USA).

### Cancer Hot Spot Panel-2 (CHP2) DNA sequencing and variant reporting

Mutations were detected by the automated Chef-ready Cancer Hotspot Panel v.2 according to the manufacturer’s protocol (Thermo Fisher Scientific, Waltham, MA, USA). Briefly, optimally 50 ng DNA (lowest 10 ng) was used as input material and loaded on to the Ion Chef Instrument. After automated library preparation, the 100 pM library pool was diluted to 35 pM, amplified on Ion Sphere Particles with Ion 510&520&530 Kit-Chef and sequenced on the Ion S5 XL using Ion 530 Chips (Thermo Fisher Scientific, Waltham, MA, USA). Variants were identified using BAM files analyzed with JSI Sequence Pilot, SeqNext V5.4.0 (JSI Medical Systems GmbH, Kippenheim, Germany) and hg19 as human reference genome. Initial screening focused on detected variants with an allele frequency of minimum 5 %. Additionally, all variants with a allele frequency lower than 5 %, but a sequencing depth of 200x were revised and documented. The pathogenicity and knowledge status of those variants were checked in publicly available databases (cBioPortal, ClinVar, dbSNP) (Cerami E *et al*., 2012; Gao J *et al*., 2013; de Bruijn et al. 2023; Landrum et al. 2018; Sherry et al. 2001). Further interpretation focused on pathogenic, likely pathogenic or variants of unknown significance, synonymous, benign or likely benign mutations were not reported.

### Organoid Culture

Human colon organoids were generated as described by Sato et. al. 2011 and Bartfeld et. al 2015. Briefly, colon endoscopy was performed using macro forceps and the biopsies were collected in ice cold basal medium containing Y(10uM). Biopsies were finely chopped, collected in a 50mL falcon tube and washed three times with 10mL of washed with ice cold PBS - (Gibco, Thermo Fisher Scientific Inc., Waltham, USA) until a clear supernatant was obtained and incubated in 2mM EDTA (Sigma) in PBS++ (Gibco, Thermo Fisher Scientific Inc., Waltham, USA) for 10 minutes at room temperature. After removal of EDTA, tissue fragments were washed with cold PBS. The fragments were then collected and added on a petridish and with the help of the glass slide, pressure was applied to release the crypts. Tissue fragments along with the released crypts were collected in 10mL ice cold basal medium and was mixed properly followed by incubation on ice for 5 min. Supernatant containing crypts was carefully collected and centrifuged at 500g for 5 min. The cell pellet was resuspended in Growth Factor Reduced Matrigel (Corning 354230) and cultured at 37 °C in the presence of 5% CO2. The organoids were then overlaid medium containing 50% CM LWRN, 20% CM R spondin, 10% CM Noggin, Primocin, Y and recombinant human EGF and 10% cocktail of factors which includes B27,N2, Nac, Nic, SB202190, A83-01.

---

**Supplementary Figure 1.**
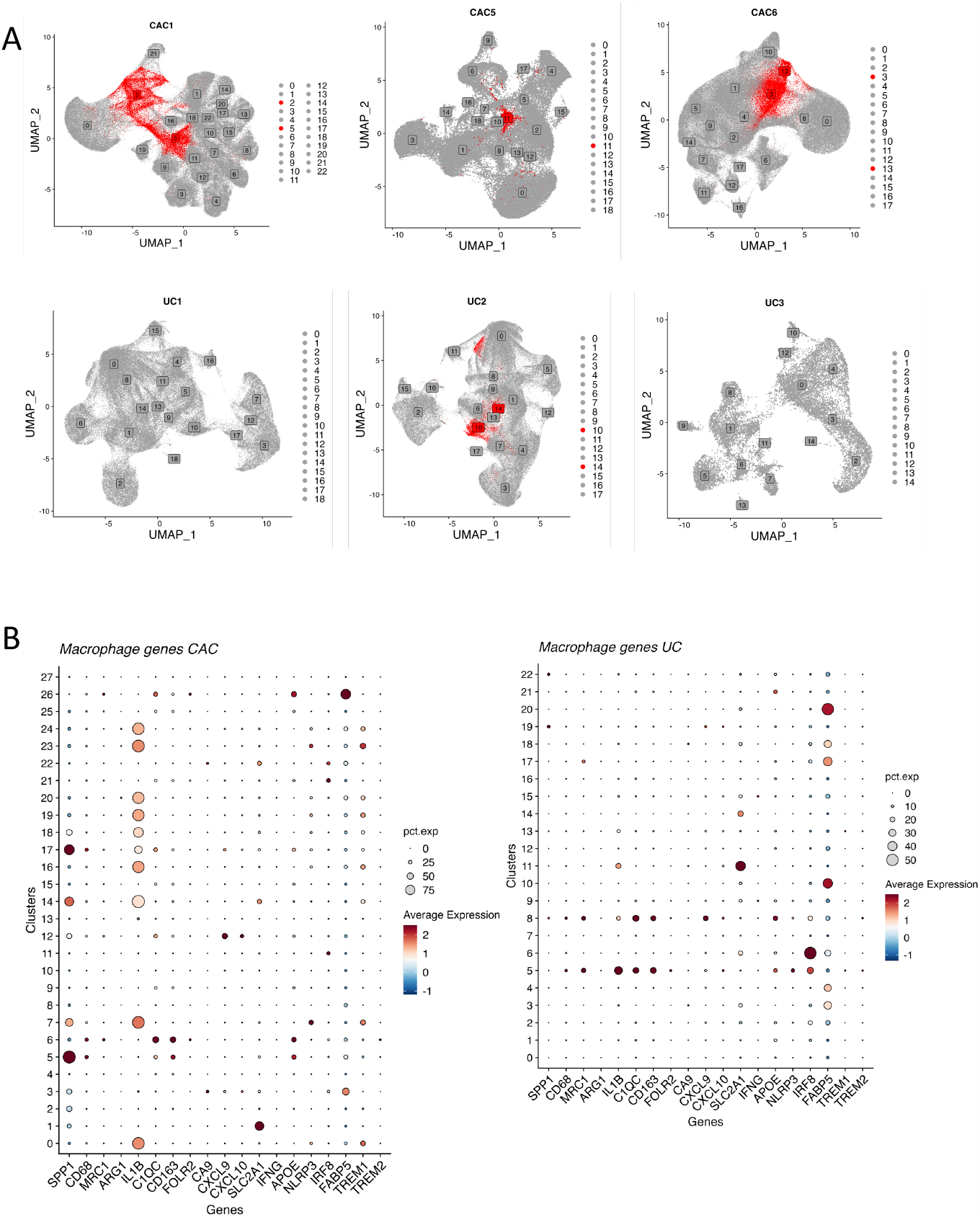
*SPP1*^*+*^ macrophage clusters are present uniformly across CAC patients. B. *SPP1*^*+*^ macrophage subset in CAC

